# deSPI: efficient classification of metagenomic reads with lightweight de Bruijn graph-based reference indexing

**DOI:** 10.1101/080200

**Authors:** Dengfeng Guan, Bo Liu, Yadong Wang

**Author notes:** The authors wish it to be known that, in their opinion, the first two authors should be regarded as Joint First Authors.

## Abstract

**Summary:** In metagenomic studies, fast and effective tools are on wide demand to implement taxonomy classification for upto billions of reads. Herein, we propose deSPI, a novel read classification method that classifies reads by recognizing and analyzing the matches between reads and reference with de Bruijn graph-based lightweight reference indexing. deSPI has faster speed with relatively small memory footprint, meanwhile, it can also achieve higher or similar sensitivity and accuracy.

**Availability:** the C++ source code of deSPI is available at https://github.com/hitbc/deSPI

**Supplementary information:** Supplementary data are available at *Bioinformatics* online.

## 1 Introduction

Metagenome sequencing is ubiquitously applied to comprehensively study environmental samples (The Human Microbiome Project Consortium, 2012; Gilbert *et al.*, 2014). In metagenomic studies, a fundamental task is to recognize the composition of the microbial community of the sequenced sample. With the ever-increasing number of sequenced genomes, it is feasible to accomplish this task by using the libraries of assembled genomes (e.g., RefSeq (Pruitt *et al.*, 2014)) as reference to implement the taxonomy classification of sequencing reads. A common approach is to align the reads against the reference (Altschul *et al.*, 1990; Huson *et al.*, 2007); however, this is not viable to handle large amount (upto terabases) of metagenomic reads due to low processing speed.

Recent efforts have been made to “pseudo-alignment” approaches (Wood and Salzburg, 2014; Ounit et al., 2015; Menzel et al., 2016; Kim et al., 2016), i.e., using short token matches between reads and reference to infer the taxonomy of the reads. In these approaches, the reference is indexed with specifically designed data structures, such as hash table or FM-index (Ferragina and Manzini, 2000) to recognize short token matches. The matches are then used for determining the taxonomy of the reads with pre-defined rules. These methods are fast. However, one of the disadvantages is that some of the methods are not space costeffective. This may be a bottleneck to integrate many genomes into reference to further improve the classification. Other methods, e.g., Kaiju (Menzel et al., 2016) and Centrifuge (Kim et al., 2016), greatly reduced the cost of memory; however, their performance could be lower than the methods using more memory, e.g., Kraken (Wood and Salzburg, 2014).

## 2 Methods

Herein, we propose deSPI, a novel short-token match-based metagenomic read classification method, which has higher speed and affordable memory cost with higher or equal sensitivity and accuracy. deSPI innovatively recognizes and analyzes the short-token matches between the reads and the reference sequences through de Bruijn graph framework. It mainly handles the reads with two key techniques.

1. Indexing: deSPI constructs the de Bruijn graph of the reference with a user-defined *k*-mer size (default: *k* = 31), and indexes the unitigs of the de Bruijn graph with a FM-index; taxonomic labels are also assigned to the indexed unitigs for further processing.
2. Classification: deSPI retrieves the maximal exact matches longer than *l* bp (default: *l* = 30) between a read and the unitigs (termed as UMEMs). The labels of the U-MEMs (which are derived from the matched unitigs) are used to infer a few paths on the taxonomy tree as evidence to classify the read.

One advantage of the method is that, with the property of de Bruijn graph, multiple *l*-mer matches having same taxonomic labels can be implicitly recognized and merged by FM-index retrieving with lower cost than that of straightforwardly retrieving all *l*-mer matches. Moreover, with the low space complexity of FM-index, the tool also has a relatively low memory footprint. A more detailed description and discussion on the deSPI method is in Supplementary Notes.

## 3 Results and conclusion

We benchmarked deSPI with two ‘pseudo’ real metagenome datasets respectively produced by Illumina HiSeq and MiSeq platforms. Each of them consists of 20 datasets (50,000 reads per dataset). Among the 20 HiSeq (MiSeq) datasets, 2(2) of them are from the species whose genomes are in RefSeq database, 14(10) of them are from the species whose genomes are homologues of RefSeq genomes, and 3 (8) of them are from the species which have at least one RefSeq genome in the same genus. Moreover, we also used the RefSeq genomes (including 1562 bacterial, 144 archaea and 36 viral genomes) to simulate 1 million Illumina-like reads with Mason simulator, to assess the ability of deSPI to handle reads from a wide variety of species.

In the benchmarking, the RefSeq genomes were used as reference. The speed, memory footprint, sensitivity and accuracy were assessed, like that of previous studies (Wood and Salzberg, 2014). Four state-ofthe-art methods, i.e., Kraken, CLARK, Kaiju and Centrifuge, were compared. All the benchmarks were conducted on a server with an Intel Xeon E4820 CPU and 1 TB RAM, running Linux Ubuntu 14.04. (More detailed information about the datasets and the implementation of the benchmarking is in Supplementary Table 1 and Supplementary Notes).

**Table 1.**
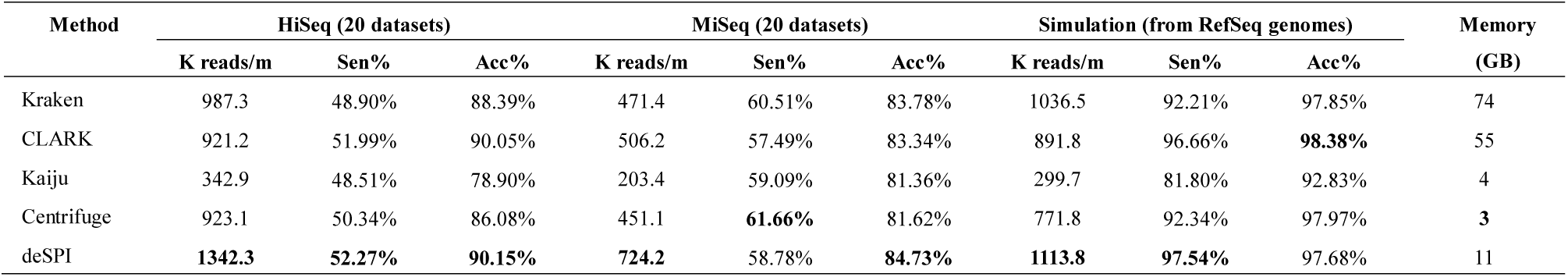
The benchmarking results. ‘HiSeq’, ‘MiSeq’ and ‘Simulation’ respectively indicate the datasets produced by Illumina platform and MiSeq platforms, and Mason simulator. ‘K reads/m’ indicates the number of thousands of reads processed per minute (in a single thread). ‘Sen%’ and ‘Acc%’ respectively indicate the sensitivity and the accuracy of the classification, which are respectively calculated as *N_R_*/*N_A_* and *N_R_*/(*N_G_* + *N_NG_*) a, where *N_R_*, *N_A_*, *N_G_* and *N_NG_* are respectively the numbers of the correctly classified reads (at genus level), all of the reads, the reads classified with the labels under or at genus level, and the resads classified with incorrect labels which are above genus level. Only the primary classification of the read is considered. The definition of sensitivity and accuracy are similar to that of the previous study (Wood and Salzberg, 2014). ‘Memory’ indicates the memory footprints (in gigabytes).

| Method | HiSeq (20 datasets) |  |  | MiSeq (20 datasets) |  |  | Simulation (from RefSeq genomes) |  |  | Memory (GB) |
| --- | --- | --- | --- | --- | --- | --- | --- | --- | --- | --- |
|  | K reads/m | Sen% | Acc% | K reads/m | Sen% | Acc% | K reads/m | Sen% | Acc% |  |
| Kraken | 987.3 | 48.90% | 88.39% | 471.4 | 60.51% | 83.78% | 1036.5 | 92.21% | 97.85% | 74 |
| CLARK | 921.2 | 51.99% | 90.05% | 506.2 | 57.49% | 83.34% | 891.8 | 96.66% | **98.38%** | 55 |
| Kaiju | 342.9 | 48.51% | 78.90% | 203.4 | 59.09% | 81.36% | 299.7 | 81.80% | 92.83% | 4 |
| Centrifuge | 923.1 | 50.34% | 86.08% | 451.1 | **61.66%** | 81.62% | 771.8 | 92.34% | 97.97% | **3** |
| deSPI | **1342.3** | **52.27%** | **90.15%** | **724.2** | 58.78% | **84.73%** | **1113.8** | **97.54%** | 97.68% | 11 |

The results suggest that deSPI has fastest processing speed. It is about 1.3 folds as fast as the tools having large memory footprints, i.e., Kraken and CLARK. Comparing to the more memory efficient tools, deSPI is about 1.5 and 3.8 folds as fast as Centrifuge and Kaiju, respectively. Meanwhile, deSPI also has affordable memory footprint with the FM-index-based reference indexing, i.e., it costs about 11 gigabytes when using RefSeq genomes as reference, which is feasible to run on most of modern PCs. Moreover, deSPI also has good scalability in parallel computing. We assessed the speed of deSPI on the HiSeq dataset with 2, 4 and 8 threads, and observed that deSPI achieved a gradually speedup (Supplementary Table 2). Comparing to Centrifuge, the advantage of deSPI on speed is more obvious with multiple threads.

deSPI and CLARK achieved best accuracies on the real sequencing and simulated datasets, respectively, indicating that reads can be handled by the two methods with less errors, while deSPI outperformed CLARK on sensitivity for all the datasets. deSPI also achieved highest sensitivities on the HiSeq and simulated datasets, indicating that it is effective to handle the reads having high sequencing quality. For the MiSeq datasets, the sensitivity of deSPI is a little lower but still comparable to the best one (Centrifuge). This is likely due to that the sequencing error rate of MiSeq reads is higher. Under this circumstance, less U-MEMs could be found with the relatively large threshold (e.g., *l* = 30). We assessed the results of deSPI with other configurations (Supplementary Table 3), and observed that, with lower thresholds (e.g., *l* = 26), the sensitivity of deSPI (60.78%) can be much closer to Centrifuge on MiSeq reads with still higher accuracy (82.97%) and speed (720.9K reads/m).

Overall, considering the speed, memory footprint, sensitivity and accuracy, deSPI can provide efficient and effective taxonomy classification for metagenomic reads. It is suited to be integrated into many metagenomics pipelines to handle large amount of reads.

## Funding

This work has been partially supported by the Nature Science Foundation of China (Nos: 61301204 and 31301089), the High-Tech Research and Development Program (863) of China (Nos: 2015AA020101, 2015AA020108 and 2014AA021505).

## Conflict of Interest

none declared.

## References

Altschul, SF, et al. (1990). Basic local alignment search tool. Journal of molecular biology, 215(3), 403–410.

Ferragina,P. and Manzini,G. (2000) Opportunistic data structures with applications. In Proceedings of the 41st Symposium on Foundations of Computer Science (FOCS 2000), IEEE Computer Society, pp. 390–398

Gilbert, JA, et al. (2014) The Earth Microbiome project: successes and aspirations. BMC Biology, 12:69

Huson, DH, etl al. (2007) MEGAN analysis of metagenomic data. Genome Res., 17(3):377–386

Kim, DS, Breitwieser, FP., & Salzberg, SL. (2016). Centrifuge: rapid and sensitive classification of metagenomic sequences. bioRxiv, 054965.

Menzel, P., Ng, K. L., & Krogh, A. (2016). Fast and sensitive taxonomic classification for metagenomics with Kaiju. Nature communications, 7.

Ounit, R., Wanamaker, S., Close, T. J., & Lonardi, S. (2015). CLARK: fast and accurate classification of metagenomic and genomic sequences using discriminative k-mers. BMC genomics, 16(1), 1.

Pruitt, KD et al. (2014) RefSeq: an update on mammalian reference sequences. Nucl. Acids Res., 42 (D1): D756–D763.

The Human Microbiome Project Consortium. (2012) A framework for human microbiome research. Nature, 486: 215–221

Wood, D. E., & Salzberg, S. L. (2014). Kraken: ultrafast metagenomic sequence classification using exact alignments. Genome biology, 15(3), 1.

